# ABCFold: easier running and comparison of AlphaFold 3, Boltz-1 and Chai-1

**DOI:** 10.1101/2025.03.12.642791

**Authors:** Luc G. Elliott, Adam J. Simpkin, Daniel J. Rigden

## Abstract

**Motivation:** The latest generation of deep learning-based structure prediction methods enable accurate modelling of most proteins and many complexes. However, preparing inputs for the locally installed software is not always straightforward, and the results of local runs are not always presented in an ideally accessible fashion. Furthermore, it is not yet clear whether the latest tools perform equivalently for all types of target.

**Results:** ABCFold facilitates the use of AlphaFold 3, Boltz-1, and Chai-1 with a standardised input to predict atomic structures, with Boltz-1 and Chai-1 being installed on runtime (if required). MSAs can be generated internally using either the JackHMMER MSA search within AlphaFold 3, or with the MMseqs2 API. Alternatively, users can provide their own custom MSAs. This therefore allows AlphaFold 3 to be installed and run without downloading the large databases needed for JackHMMER. There are also straightforward options to use templates, including custom templates. Results from all packages are treated in a unified fashion, enabling easy comparison of results from different methods. A variety of visualisation options are available which include information on steric clashes.

**Availability and implementation:** ABCFold is coded in Python and JavaScript. All scripts and associated documentation are available from https://github.com/rigdenlab/ABCFold or https://pypi.org/project/ABCFold/1.0.0/.

## 1 Introduction

The latest generation of deep learning-based methods, such as AlphaFold 3 (Abramson et al., 2024), Boltz-1 (Wohlwend et al., 2024) and Chai-1 (Discovery *et al*., 2024), allow for the accurate modelling of proteins, nucleic acids, complexes and ligands. Some comparative exercises have been published (Morehead et al., 2024; Škrinjar *et al*., 2025; Medvedev et al., 2025) but it remains largely unclear what idiosyncratic strengths and weaknesses these methods - like previous generations (Pratt et al., 2025; Wojciechowska *et al*., 2024) - may have.

ABCFold is designed to make it easier to run AlphaFold 3, Boltz-1 and Chai-1. Each method takes in a unique input for defining advanced features such as ligands, post-translational modifications, and distance restraints. ABCFold simplifies operation by taking a single input in the format of an AlphaFold 3 JSON and automatically converting it to work with Boltz-1 and Chai-1. Additionally, ABCFold makes it straightforward to run these methods with custom Multiple Sequence Alignments (MSA), with the popular MMseqs2 API (Steinegger and Söding, 2017; Mirdita *et al*., 2019) employed for MSA generation and template selection (AlphaFold 3 and Chai-1), and with custom templates (AlphaFold 3 only). It also remaps and reorders output chains resulting from distinct treatments by the three programs, further facilitating comparison.

Equally importantly, ABCFold provides an information-rich comparative output that allows users to see differences in their results. Models from all three methods can be ranked according to a variety of scores including number of clashes. A feature-viewer (Paladin *et al*., 2020) provides zoom-able side-by-side comparison of colour-coded pLDDT values. And finally PAE viewer (Elfmann and Stülke, 2023) is used for interactive exploration of the Predicted Aligned Error (PAE) and molecular coordinates, with clashes marked in both contexts. Thus, while not the only package aiming to aid running the latest methods for structure prediction (Ford *et al*., 2025), the visualisation and comparison options of ABCFold may help the community reach a quicker assessment of the new methods’ strengths and weaknesses.

## 2. Results

### 2.1 Simplifying inputs

With ABCFold, users only need to prepare a single input JSON file in the format of AlphaFold 3, to run AlphaFold 3, Boltz-1 and Chai-1. ABCFold can be run using: `abcfold <input_json> <output_dir> -abc` in its simplest form, wherein the latter flags, in this case, trigger running of all three methods according to their initial. The input JSON is automatically converted into the correct format for Boltz-1 and Chai-1, allowing the methods to be run sequentially from a single input.

### 2.2 MSAs and templates

MSA selection plays an important role in the performance of deep learning based prediction methods (Jumper *et al*., 2021; Simpkin et al., 2023). It is therefore useful to be able to incorporate custom MSAs and templates into such methods. An added benefit when using MMseqs2 and ABCFold is the removal of the need for a local database search for MSA and template generation: thus, users can avoid having to download the large databases locally on their systems. If running AlphaFold 3 with the native JackHMMER search, the resulting MSA will automatically be used for Boltz-1 and Chai-1 runs.

#### 2.2.1 MMseqs2 MSA

ABCFold has been designed to facilitate the use of the MMseqs2 API in AlphaFold 3. Specifying the `-mmseqs2` command line flag will leverage the ColabFold MMseqs2 API (Mirdita et al., 2022; Steinegger and Söding, 2017) to perform an MSA search against UniRef30 (v2303) and colabfold_envdb (v202108). The resulting MSA is passed into AlphaFold 3, Boltz-1 and Chai-1, thus avoiding the computational overhead of recalculating it.

#### 2.2.2 Custom MSAs

As in AlphaFold 3, ABCFold can be run with a custom MSA by modifying the input JSON to contain a path to the custom MSA using the `unpairedMsaPath` or `unpairedMSA` variable:

~~~
{
 “protein”: {
       “id”: “A”,
       “sequence”: “PVLSCGEWQL”,
       “modifications”: [
       {“ptmType”: “HY3”, “ptmPosition”: 1},
       {“ptmType”: “P1L”, “ptmPosition”: 5}
       ],
       “unpairedMsa”: …,
       “unpairedMsaPath”: …,
       “templates”: […]
 }
}
~~~

This MSA path specified at `unpairedMsaPath`, or the MSA string at `unpairedMsa`, will be passed into AlphaFold 3, Boltz-1 and Chai-1, allowing a fair comparison of the methods.

#### 2.2.3 Templates

AlphaFold 3 will routinely use templates, and as of version 0.6.0, Chai-1 can also use templates if Kalign (Lassmann, 2019) is installed. ABCFold allows users to run AlphaFold 3 and Chai-1 with or without templates, and also provides the option of using templates found by MMSeqs2 or the templates found by AlphaFold 3. The use of templates has been shown to significantly improve the quality of the output models in some instances (Keegan et al., 2024) and it can therefore be important to use templates when they are available.

Similarly, users may wish to input custom templates (currently only supported by AlphaFold 3) using the `--custom_template` flag and providing structure file paths. If using a multi-chain template, they must also specify which chain to use with the `-- custom_template_chain` flag, and if modelling a complex, they must specify the target ID the template corresponds to using the `--target_id` flag. An example of a more complex ABCFold input where custom templates are used with MMSeqs2 whilst only running Alphafold is as follows:

~~~
abcfold <input_json> <output_dir> -a –
custom_template <path_to_pdb_or_cif> --
custom_template_chain A --target_id B --mmseqs2
~~~

A full list of flags can be found on the ABCFold git repository: https://github.com/rigdenlab/ABCFold or the PyPi project page: https://pypi.org/project/ABCFold/1.0.0/.

### 2.3 AlphaFold 3, Boltz-1 and Chai-1 installation and version checks

Where the `-b / --boltz1` or `-c / --chai1` flags are specified, ABCFold will, if necessary, install Boltz-1 and Chai-1 before running them. It is possible for the user to install Boltz-1 and Chai-1 before running but ABCFold will install the corresponding version of each software that is compatible with ABCFold. At the time of writing, this is `boltz==0.4.1` and `chailab==0.6.0`.

AlphaFold 3 can be installed from the GitHub page (https://github.com/google-deepmind/alphafold3). At time of writing ABCFold supports versions >= 3.0.0. Users must also obtain model parameters for AlphaFold 3, subject to their terms and conditions, but database installation is only required if the user wishes to use the native AlphaFold 3 JackHMMER-based MSA search.

### 2.4 Treatment of results

When running ABCFold, AlphaFold 3, Boltz-1 and Chai-1 each give, by default, five output structure predictions, but this can be increased by using the `--number_of_models` flag. For Chai-1, the submission script is modified to extract the PAE values for the output models. Results from all specified runs are made available in the user-specified output directory, a simplified sub-directory structure of the output directory is shown below for the example 8aq6.

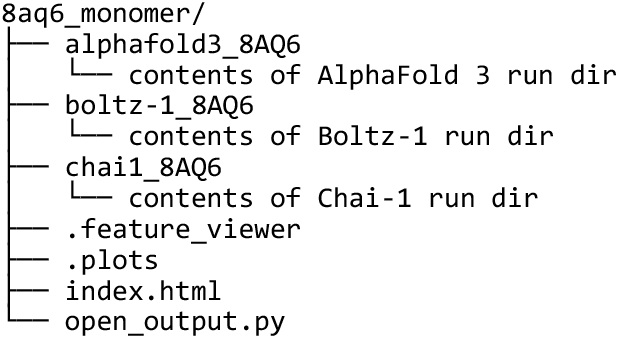

Boltz-1 and Chai-1 both label the input chains in their output with consecutive uppercase Alphabet characters, i.e ABC… AlphaFold 3 maintains the same sequence id mapping as the input JSON, but orders the chains according to label. For consistency of output between AlphaFold 3, Boltz-1 and Chai-1, the chain labels of their output models are remapped to the label ids given in the input JSON.

### 2.5 Visualisation of results

ABCFold has been designed with the goal of comparing models in mind. The main output page (Fig 1a) shows a table of results ranked by average pLDDT. Also shown are scores such as H-score (Simpkin et al., 2022), and the number of atomic clashes are defined as shown below:

**Fig. 1.**
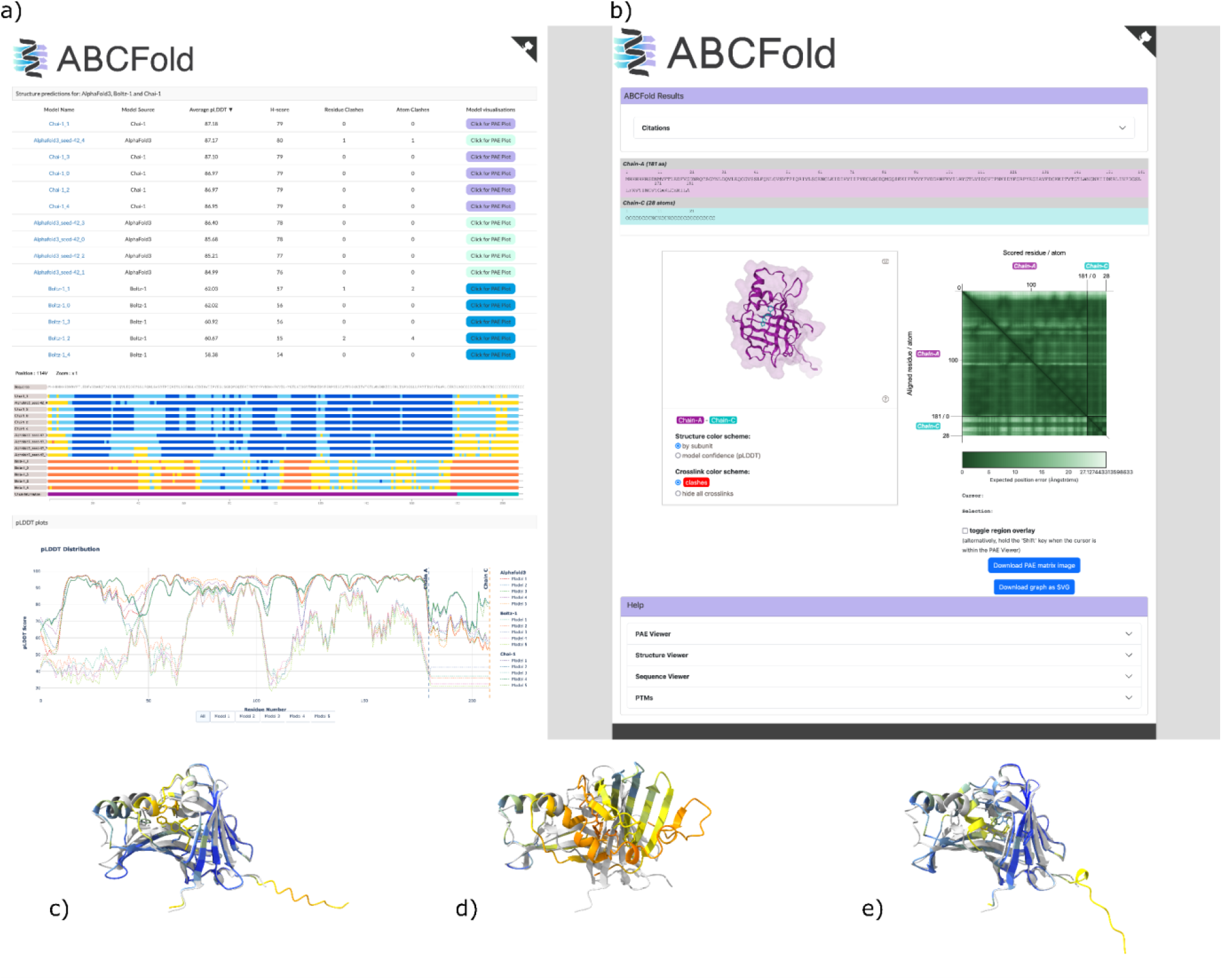
Visualisation of results in ABCFold. a) The main ABCFold output page for 8aq6. This shows the output table, the feature-viewer representation of pLDDT and the Plotly pLDDT plot and allows comparison between the different methods. b) The PAEViewer output page shows a 3D representation of the model, a PAE figure, and the location of possible clashes. In c) to e) are superpositions of the crystal structure (grey) with the best model from each of the methods (coloured by pLDDT) created with ChimeraX (Pettersen et al., 2021). In this example, which should in no way be considered as representative of relative performance in a broader sense, the best models from c) AlphaFold 3, d) Boltz-1 and e) Chai-1 have TM-scores (Zhang and Skolnick, 2004) of 0.90, 0.45 and 0.87, respectively.

1. 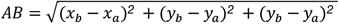
2. 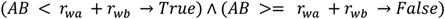

Where AB is distance between the xyz Cartesian coordinates of atom a and b, respectively (*x*_*a*_, *y*_*a*_, *z*_*a*_), (*x*_*b*_, *y*_*b*_, *z*_*b*_). *r*_*w*_ represents the van der Waals radius of an atom. True represents a clash between the atoms. Also shown on the output page (Fig1A) are the number of predicted residue clashes i.e. the number of residues in a model within which at least one atom is clashing.

A visual representation of the pLDDT scores is shown using feature-viewer (Paladin *et al*., 2020), where very low confidence regions (pLDDT<50) are shown in orange, low confidence regions (50 < pLDDT < 70) are shown in yellow, confident regions (70 < pLDDT < 90) are shown in light blue and very confident regions (pLDDT > 90) are shown in dark blue. Below this is an interactive pLDDT plot made with Plotly (GitHub - plotly/plotly.py: The interactive graphing library for Python :sparkles:) that allows for a finer-grained comparison of the methods.

By default, none of the three methods provide PAE plots. ABCFold will generate them for each model using PAEViewer (Elfmann and Stülke, 2023) (Fig 1b). Residues involved in clashes are indicated with red circles in the PAE, but the clash display can be switched off. The NGLviewer (Rose and Hildebrand, 2015) panel allows interactive exploration of the coordinates and their confidence, linked to the PAE, and indicates clashes as red bars between Calpha atoms by repurposing the inter/intra protein link facility.

ABCFold creates a server locally to view the output page(s). If working on a cluster without port forwarding / tunneling available, the server would not be accessible. ABCFold includes the flags `- -no_server` to stop the server starting automatically; the server can then be started on a local system with the provided `open_output.py` in the output directory. Alternatively no html pages will be generated if the flag `--no_server` is provided.

## Data Availability

The source code, installation instructions and input examples can be located at https://pypi.org/project/ABCFold/1.0.0/ or https://github.com/rigdenlab/ABCFold. ABCFold should ideally be installed directly through pip: `pip install ABCFold` or directly from source from the git repository.

## Funding

This work was supported by the Biotechnology and Biological Sciences Research Council (BBSRC), Project BB/Y008901/1

## Conflict of Interest

none declared.

## Notes

### Competing Interest Statement

The authors have declared no competing interest.

### Summary of Updates

Formatting errors in the equation and directory structure elements have been corrected

